## Supplementary Figures for "Uncovering circuit mechanisms of current sinks and sources with biophysical simulations of primary visual cortex"

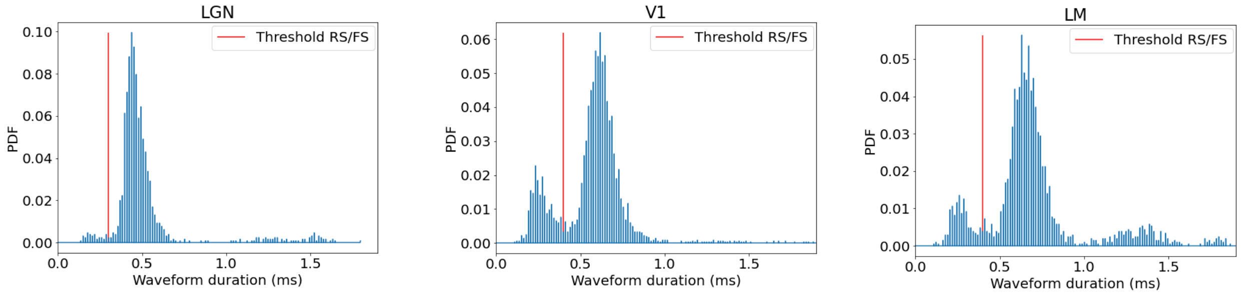


**Figure S1:** Distributions of waveform duration in cells from LGN, V1, and LM and the threshold (red line) between classifying as regular-spiking (RS) or fast-spiking (FS).

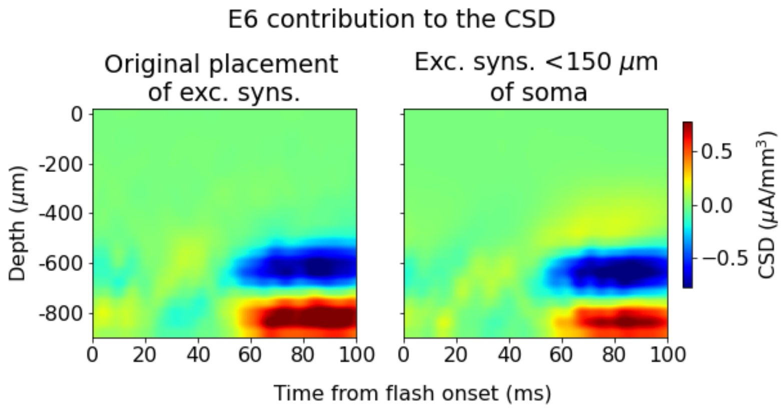


**Figure S2:** Contributions to the total CSD from L6 excitatory cells with the original placement of recurrent excitatory synapses uniformly along the whole length of their dendrites (left) and after moving all recurrent excitatory synapses within 150 μm from the soma.

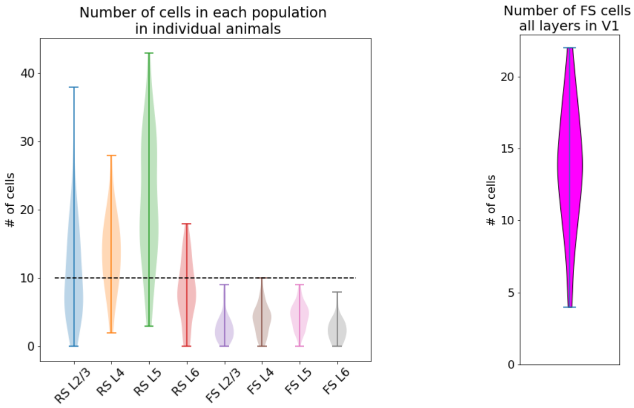


**Figure S3:** Left: Number of regular-spiking (RS) and fast-spiking (FS) in each layer in individual animals. Right: Number of FS cells across all layers in V1 in individual animals.

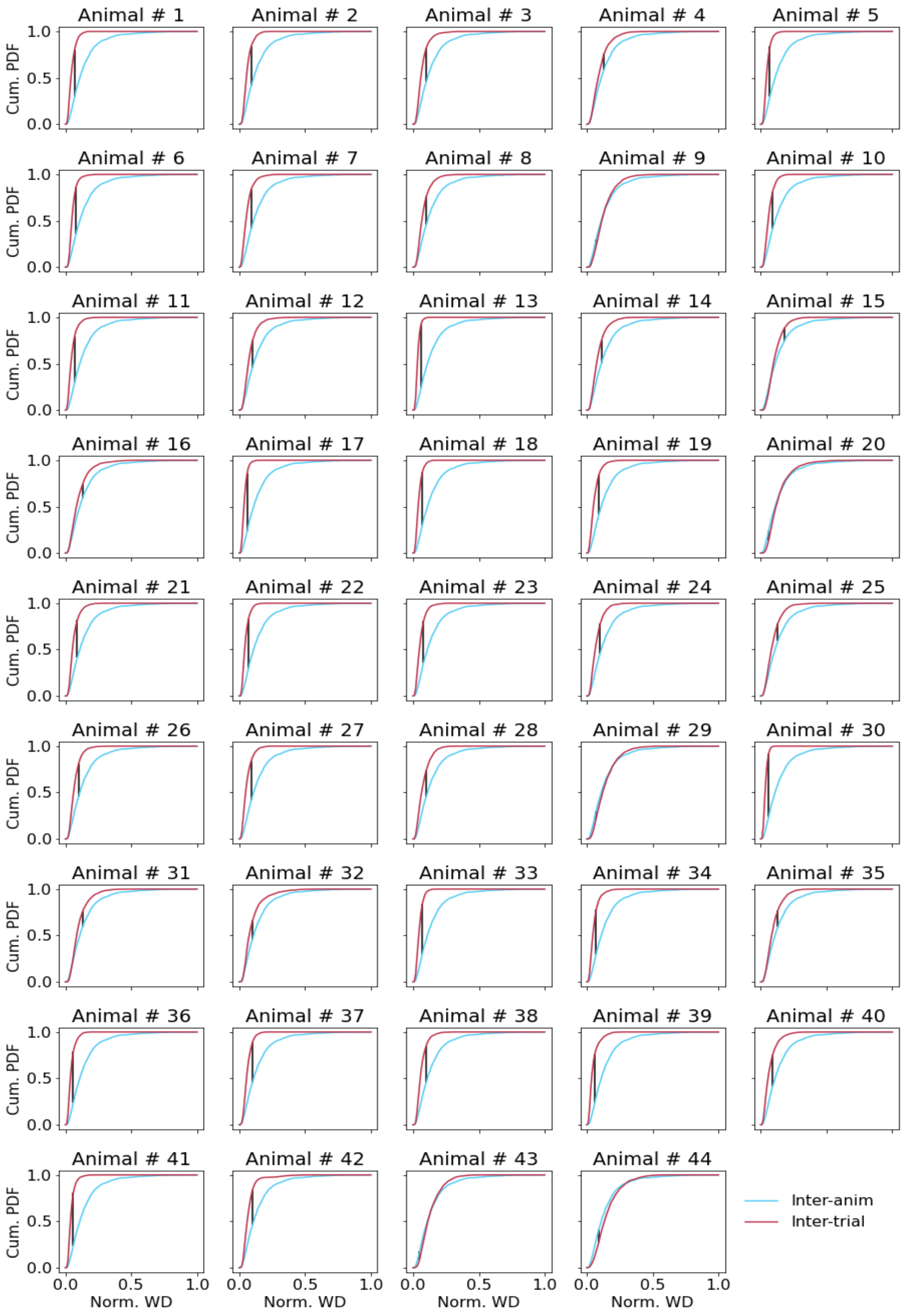


**Figure S4:** Cumulative distributions of pairwise WDs between trial-averaged CSD of individual animals (blue line) and pairwise WDs between single trial CSD in each animal (red lines). The black line denotes the point of maximal distance between the two distributions.

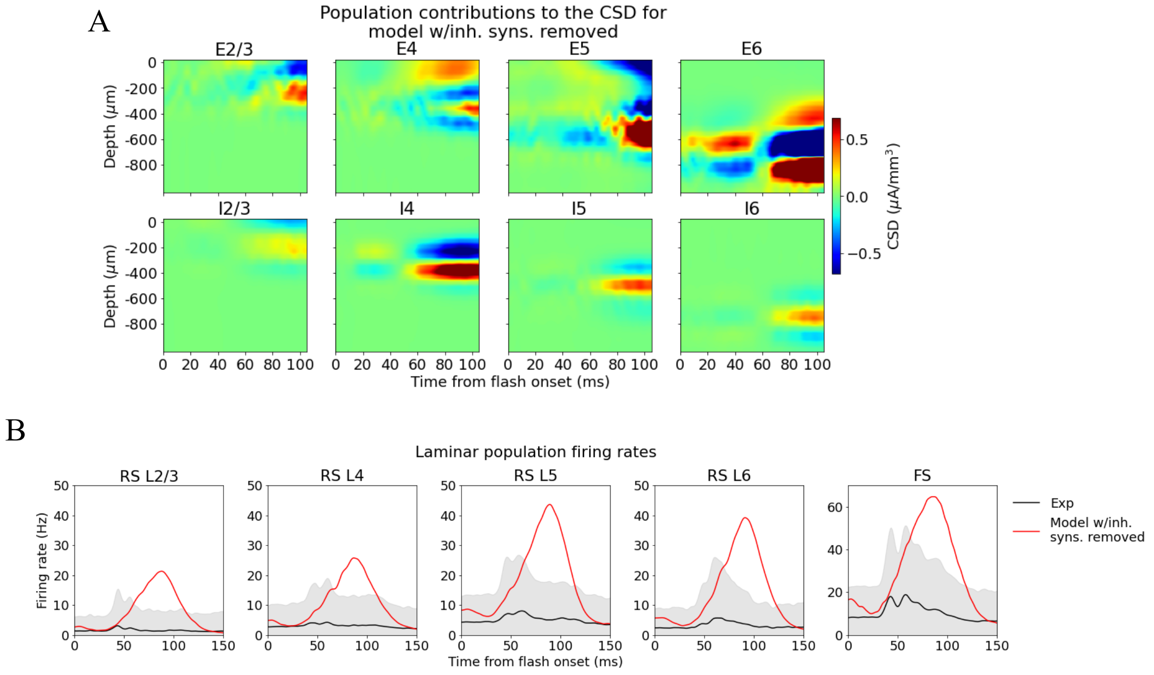


**Figure S5: (A)** Population contributions to the total CSD and **(B)** laminar population firing rates in a simulation where all inhibitory synapses have been removed.

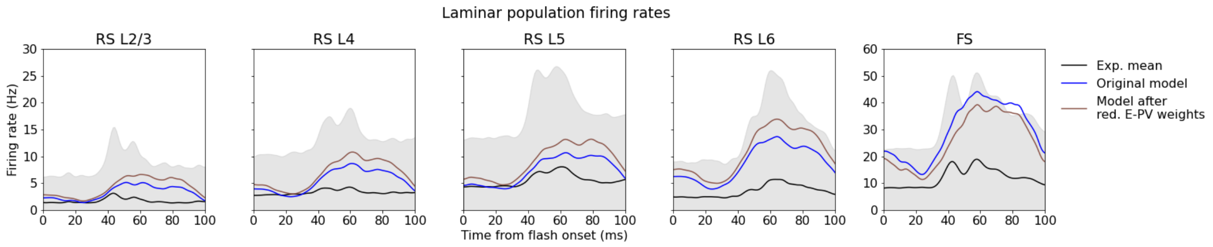


**Figure S6:** Laminar population firing rates in original model (blue line), and in the model after reduction of recurrent excitatory synaptic weights to all Pvalb cells by 30 % (brown line) and in experiments (black line).

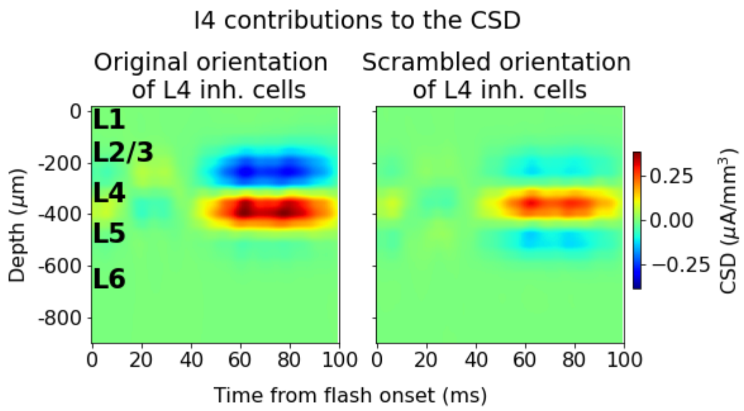


**Figure S7:** CSD contribution of L4 inhibitory cells with original orientation (left) and scrambled orientation (right).
